# Myocardial Elasticity Imaging Correlates with Histopathology in a Model of Anthracycline-Induced Cardiotoxicity

**DOI:** 10.1101/2023.10.11.561881

**Authors:** Caroline E. Miller, Jennifer H. Jordan, Alexandra Thomas, Sarah R. Friday, Giselle C. Meléndez, Jared A. Weis

## Abstract

**Background:** There is considerable focus on developing strategies for identifying subclinical cardiac decline prior to cardiac failure. Myocardial tissue elasticity changes may precede irreversible cardiac damage, providing promise for an early biomarker for cardiac decline. Biomarker strategies are of particular interest in cardio-oncology due to cardiotoxic effects of anti-neoplastic therapies, particularly anthracycline-based chemotherapeutics. Current clinical methods for diagnosing cardiotoxicity are too coarse to identify cardiac decline early enough for meaningful therapeutic intervention, or too cumbersome for clinical implementation.

**Methods:** Utilizing changes in myocardial elasticity as a biomarker for subclinical cardiac decline, we developed a biomechanical model-based elasticity imaging methodology (BEIM) to estimate spatial maps of left ventricle (LV) myocardial elasticity. In this study, we employ this methodology to assess changes in LV elasticity in a non-human primate model of doxorubicin-induced cardiotoxicity. Cardiac magnetic resonance imaging of five African Green monkeys was acquired at baseline prior to doxorubicin administration, 6-weeks, and 15-weeks after final doxorubicin dose and histopathological samples of the LV were taken at 15-weeks after final doxorubicin dose. Spatial elasticity maps of the mid-short axis plane of the LV were estimated at each image acquisition. Global and regional LV elasticity were calculated and changes between imaging time points was assessed. LV elasticity at baseline and final time point were compared to cardiomyocyte size and collagen volume fraction measurements calculated from histopathological staining of archived tissue bank samples and study endpoint tissue samples utilizing Pearson’s correlation coefficients.

**Results:** We identify significant changes in LV elasticity between each imaging time point both globally and regionally. We also demonstrate strong correlation between LV elasticity and cardiomyocyte size and collagen volume fraction measurements. Results indicate that LV elasticity estimates calculated using BEIM correlate with histopathological changes that occur due to doxorubicin administration, validating LV elasticity solutions and providing significant promise for use of BEIM to non-invasively elucidate cardiac injury.

**Conclusions:** This methodology can show progressive changes in LV elasticity and has potential to be a more sensitive indicator of elasticity changes than current clinical measures of cardiotoxicity. LV elasticity may provide a valuable biomarker for cardiotoxic effects of anthracycline-based chemotherapeutics and cardiac disease detection.

## Introduction

The early detection of cardiotoxicity induced by chemotherapy treatment in cancer patients has become increasingly important^1, 2^ since left ventricular (LV) deterioration of ejection fraction (EF) indicates significant, and often irreversible adverse myocardial remodeling^3, 4^. Cardiac magnetic resonance (CMR)-derived global circumferential strain (GCS) by feature tracking strain conducted after initiation of cardiotoxic chemotherapy is prognostic for the development of cancer therapy–related cardiac dysfunction (CTRCD), especially in women with breast cancer receiving anthracyclines and trastuzumab^5, 6^.

Strain imaging is based on the analysis of the gradients of ventricular deformation occurring within a cardiac cycle as a combined indicator of LV contractility and stiffness^7–9^. While strain is related to tissue elasticity, it is not a direct intrinsic measurement of tissue elasticity. The surrogacy of strain imaging as a sensitive metric of tissue elasticity relies on the problematic assumption of uniform mechanical stress within the ventricle, however, this assumption does not hold true due to variability of the cardiac geometry and myocardial fiber throughout the cardiac cycle^10^. Moreover, strain imaging analysis is also confounded by changes in ventricle shape, size, and pre-loading (hypovolemia), which are common among patients undergoing anthracycline-based chemotherapies^11, 12^. These confounding factors can result in observations of apparent strain imaging deterioration during treatment and/or strain imaging recovery after discontinuation of therapy that may be biased passive indicators of changes in geometry and hypovolemia and not reflective of true cardiac tissue health^13^.

Based on the direct relationship between myocardial stiffness and cardiac function, tissue elasticity may be an additional important biomarker of cardiotoxicity that can be derived from CMR. However, there remains a need for the development of a validated non-invasive imaging methodology to directly measure myocardial elasticity prior to its utilization in future studies to determine if it may be an early predictor. To address this need, we developed a non-invasive biomechanical model-based elasticity imaging methodology (BEIM) to estimate LV elasticity utilizing routinely acquired cine CMR images^14^ and have characterized the methodological reproducibility^15^. Briefly, BEIM employs an inverse biomechanics computational modeling framework to estimate a direct intrinsic measurement of LV tissue elasticity that is not dependent upon uniform stress assumptions and not confounded by changes in LV shape or size. In this study, we utilized BEIM to estimate LV elasticity in a non-human primate anthracycline-induced cardiotoxicity model that employs a human-equivalent doxorubicin (Dox) dosing protocol that reliably induces severe LV dysfunction. Importantly, at the histopathological level, this animal model exhibits extensive interstitial cardiac fibrosis^16^. The goal of our study is to validate BEIM for assessing spatiotemporal changes in LV elasticity at baseline and after Dox administration using routine CMR imaging.

## Materials and Methods

### Animals

This study conformed to the principles of the National Institutes of Health and all protocols were approved by Wake Forest University (WFU) Animal Care and Use Committee. We conducted BEIM elasticity measurements in repository longitudinal CMR images acquired from a cohort of five African Green monkeys (AGM) treated with human-equivalent doses of Dox based on an allometric scaling of body surface area (BSA)^16^. Animals (aged 13 ± 1.3 years) underwent Dox treatment consisting of 2 doses of 30 mg/m^2^ and 3 doses of 60 mg/m^2^ administered biweekly, IV (total cumulative dose of 240 mg/m^2^) ^16^. Animals underwent CMR imaging before, 6-weeks, and 15-weeks after the last dose of Dox. After euthanasia was induced, hearts were excised and a transverse mid-section corresponding to the LV mid-short axis plane from CMR was fixed in 4% paraformaldehyde for subsequent histopathology analysis. For non-treated control samples, archival tissue-banked tissues from five age- and gender-matched healthy animals were used for histopathological assessment comparisons due to ethical considerations.

### Collagen Volume Fraction and Cardiomyocyte Numbers and Size

As previously described^16^, five-micron sections from each block were stained with collagen specific picrosirius red (0.1% Sirius Red F3BA in picric acid) for collagen volume fraction quantification (CVF)^17–19^. Twenty random fields were acquired at 20X, excluding perivascular collagen, for collagen volume fraction quantification using Image J (NIH, Bethesda, MA) and expressed as a percentage of area. Cardiomyocyte area and cardiomyocyte number were quantified by identifying cell membranes through staining tissue samples with fluorescent-tagged wheat germ agglutinin (WGA) and quantified using Mantra™ quantitative pathology workstation (PerKinElmer, Waltham, MA)^17^. Quantification of cell size was done using inForm® image analysis software (PerKinElmer, Waltham, MA).

### CMR Imaging and Image Processing

CMR imaging was performed on a 3T Siemens Skyra Scanner before (baseline), 6 and 15 weeks after the final dose of Dox. A cardiac short-axis cine steady-state free procession (SSFP) sequence with a 192x128 matrix, 20cm field of view, 10ms repetition time, 4ms echo time, a 35-degree flip angle, 40ms temporal resolution, and 6mm slice resolution was utilized. The LV wall was segmented from previously acquired images using AI-based segmentation software Segment Medviso^20^, and used to calculate LVEF based on Simpson’s rule^21, 22^. Offline image contours were performed by an experienced observer (GCM). Global and segmental peak circumferential strain was calculated for each image acquisition in the short-axis orientation utilizing feature tracking over an entire cardiac cycle with the available software Segment v3.0^20^. Cardiac output values were calculated as the product of stroke volume and heart rate. For each image, time frames of the cardiac cycle were automatically selected to represent passive diastole of the LV by assessing LV volume measurements throughout the cardiac cycle calculated using Segment Medviso. The mid time frame of passive diastole was then automatically selected and used as a reference for non-rigid diffeomorphic demons registration of each time frame to estimate passive deformation of the LV with a standard deviation of 25 and 3 resolution levels^23–25^. Gaussian smoothing with a standard deviation of 4 was applied to the resulting passive deformation fields. A finite element mesh was generated from LV wall segmentations with a nominal edge length of 0.6 mm and clustered into 150 regions for property reconstruction with an average region size of 2 mm^2^ using k-means clustering based on Euclidean distances.

### Biomechanical Model-based Elasticity Imaging Methodology (BEIM)

We utilize a previously developed biomechanical model-based elasticity imaging methodology (BEIM)^14^ to estimate spatial elasticity maps of the LV in AGMs. While a detailed description of the method can be found in our prior work^14, 15^, a brief description follows. Observed deformation representing inherent cardiac motion during passive diastole was obtained from non-rigid image registration. Deformation at the LV boundaries was extracted and applied to the finite element mesh as boundary conditions for subsequent elasticity reconstruction. Model estimated deformation was calculated based on a spatial elasticity distribution input into a transversely isotropic linear elastic biomechanics model and solved over the finite element mesh using the Galerkin method of weighted residuals. Poisson’s ratio was assumed to be nearly incompressible at 0.45 in the circumferential and radial directions to decrease degrees of freedom for solving the biomechanics model. Model estimated deformation was compared to observed deformation in a region-based inverse elasticity parameter estimation framework. Properties within each region representative of circumferential and radial elastic modulus were iteratively updated based on an L-BFGS quasi-Newton algorithm with parameter sensitivity gradients calculated using the adjoint method^26, 27^. The objective function, defined as the sum squared error between observed and model-estimated deformation, was minimized until a preset convergence tolerance was met. The result of the method is spatial circumferential and radial elastic modulus maps of the LV in the circumferential and radial directions. As described in previous work^15^, since fully-constrained displacement-based boundary condition solutions are indeterminate, we normalize elasticity solutions by a time-stress integral calculated over the endocardium^28^ with the time-stress-integral serving as a normalization factor to allow for image comparisons across subjects and time points and representative of the stress on the endocardium throughout passive diastole based on the spatial elasticity solution. Normalized circumferential, radial, and shear modulus maps were then compared across image time point acquisitions. A global average was used for subsequent comparisons across image acquisitions, and segment averages were assessed to compare spatially dependent changes in modulus. For segmental analysis, the LV was divided into three anatomical regions that represent septal, inferior, and anterior segments of the LV, corresponding to the LV pathological segments. Segment locations were defined by selecting a point on the LV that corresponds to the mid-point of the right ventricle. Based on LV point selection, three equal anterior segments and three inferior segments were defined to create a six anatomical segment model representative of the American Heart Association (AHA) LV model for short-axis mid-cavity slices^29^. To assess region-based changes that correspond to collected pathological tissue samples, the two septal segments, two posterior segments, and two anterior segments were each combined to create septal, posterior, and anterior segments, respectively. Note that previous work shows minimal intraobserver differences in BIEM-calculated elasticity^15^.

### Statistical Analysis

Biometric and cardiac function metrics were calculated from each CMR image to indicate cardiotoxicity in the AGM model of anthracycline-induced cardiotoxicity and compared using one-way ANOVA. LV histopathology was assessed by comparing cardiomyocyte area and CVF values of the age-matched tissue bank samples to necropsy samples of the AGM model acquired at 15-weeks using an unpaired t-test. Both analyses were previously reported for this cohort^16^. Regional LV circumferential, radial, and shear elasticity maps for each subject were estimated based on mid-axis cine cardiac MR images acquired at baseline, 6-weeks, and 15-weeks. Average global modulus values and segment-based values were compared across each image acquisition using paired t-tests. Changes in elasticity between image acquisitions and regions were assessed using two-way ANOVA. For correlation studies, baseline elasticity values were compared to control tissue bank samples of age-matched AGM subjects and elasticity values calculated from images acquired at 15-weeks were compared to tissue samples of the AGM model acquired at 15-weeks. Correlations between average global modulus values and CVF and cardiomyocyte area were assessed using Pearson’s correlation coefficients, R^2^ and corresponding *p*-values. Peak global and segmental circumferential strain was calculated at corresponding mid-axis locations for each CMR image acquisition and segment using Segment software (Medviso)^20, 30, 31^ and compared across image time points using paired t-tests and across segments and time points utilizing two-way ANOVA. Strain correlations were also assessed by comparing strain to CVF and cardiomyocyte area using Pearson’s correlation. All analyses were completed using GraphPad Software and a *p*-value of <0.05 was considered significant.

## Results

### Biometrics and CMR Parameters

Sample biometrics and cardiac function measurements were calculated from each imaging acquisition to confirm cardiotoxicity due to Dox and shown in Table 1. Sample body surface area and LV mass index significantly differed between image acquisitions indicated by a *p*-value of 0.025 and 0.024 respectively. LVEF and CO measurements indicate significant cardiotoxicity with an average decrease of 25% and 1,271 ml/min, respectively, representing 35% and 52% decreases. LVEF and CO significantly differed between imaging time points indicated by a *p*-value of 0.012 and 0.0012 respectively. LV end diastolic and end systolic volumes were also assessed for each image acquisition but did not significantly differ between timepoints.

**Table 1:**
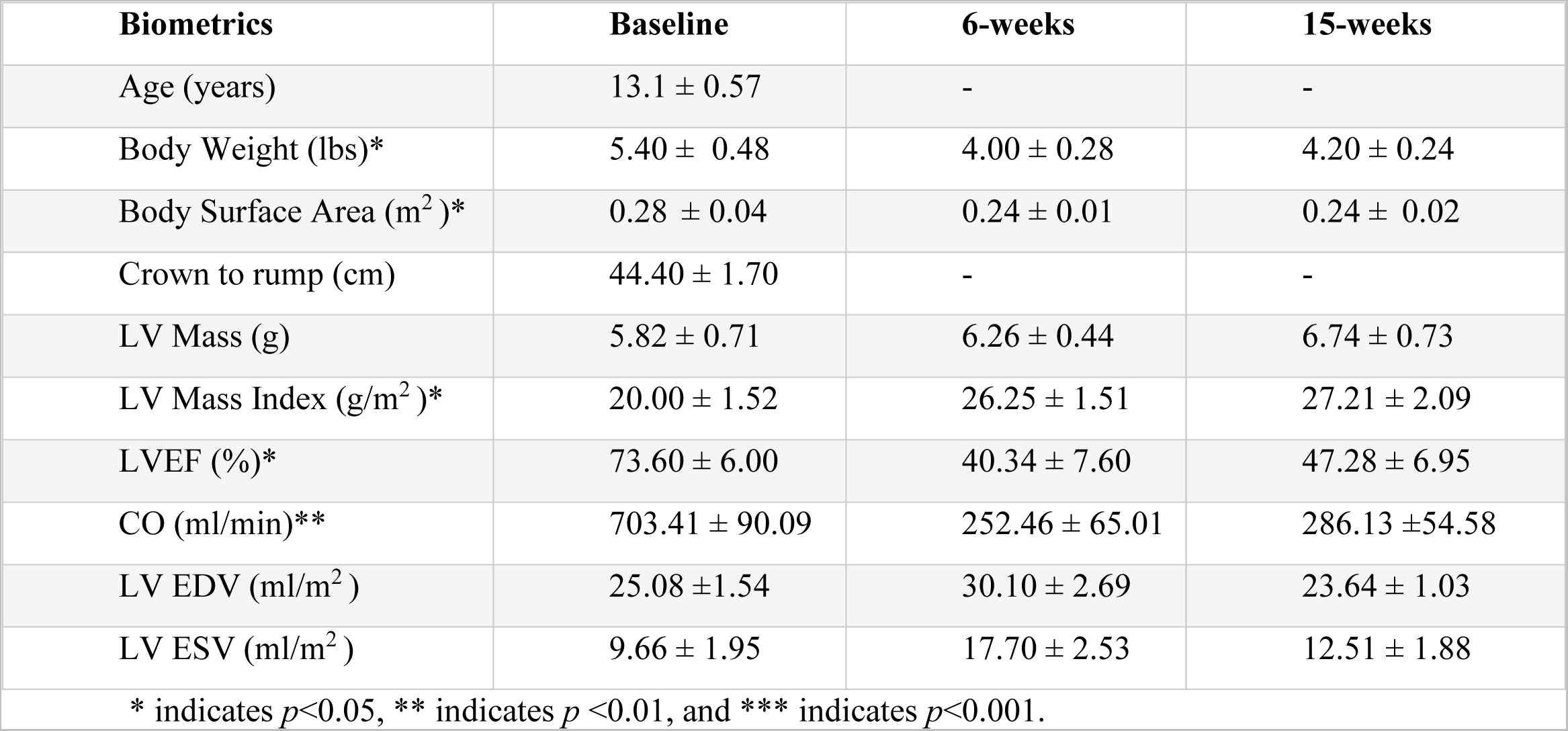
Biometric and Cardiac MR Imaging Parameters for AGM Subjects (Average ± Standard Error.

| **Biometrics** | **Baseline** | **6-weeks** | **15-weeks** |
| --- | --- | --- | --- |
| Age (years) | 13.1 ± 0.57 | - | - |
| Body Weight (lbs)* | 5.40 ± 0.48 | 4.00 ± 0.28 | 4.20 ± 0.24 |
| Body Surface Area (m<sup>2</sup>)* | 0.28 ± 0.04 | 0.24 ± 0.01 | 0.24 ± 0.02 |
| Crown to rump (cm) | 44.40 ± 1.70 | - | - |
| LV Mass (g) | 5.82 ± 0.71 | 6.26 ± 0.44 | 6.74 ± 0.73 |
| LV Mass Index (g/m<sup>2</sup>)* | 20.00 ± 1.52 | 26.25 ± 1.51 | 27.21 ± 2.09 |
| LVEF (%)* | 73.60 ± 6.00 | 40.34 ± 7.60 | 47.28 ± 6.95 |
| CO (ml/min)** | 703.41 ± 90.09 | 252.46 ± 65.01 | 286.13 ± 54.58 |
| LV EDV (ml/m<sup>2</sup>) | 25.08 ± 1.54 | 30.10 ± 2.69 | 23.64 ± 1.03 |
| LV ESV (ml/m<sup>2</sup>) | 9.66 ± 1.95 | 17.70 ± 2.53 | 12.51 ± 1.88 |

\* indicates $p < 0.05$ , \*\* indicates $p < 0.01$ , and \*\*\* indicates $p < 0.001$ .

### LV Elasticity Changes with Doxorubicin Administration

Circumferential, radial, and shear modulus maps were estimated using BEIM from imaging data acquired at baseline, 6-weeks, and 15-weeks after doxorubicin, shown in Figure 1. Average global elastic modulus values were calculated from each image to quantify changes in modulus over time. Average modulus at each image acquisition was statistically compared using paired t-tests with statistical significance indicated by asterisks. As shown in Figure 1A-C, we show significant increases in modulus between each image acquisition. We observe a significant progressive increase in modulus over time as a result of doxorubicin administration. Progressive spatial stiffening is also demonstrated in representative elasticity maps for each modulus at each image acquisition for all subjects and is shown in representative images of one subject in Figure 1D-F. To indicate changes in CVF, histopathological samples at 15-weeks following final Dox dose and histopathological controls obtained from tissue bank samples with comparable age and no indications of cardiovascular disease were both stained with picrosirius red to indicate collagen deposition. Figure 1G-J shows significant increases in collagen throughout the entire ventricle from the control samples to samples acquired 15-weeks after the final Dox dose.

**Figure 1:**
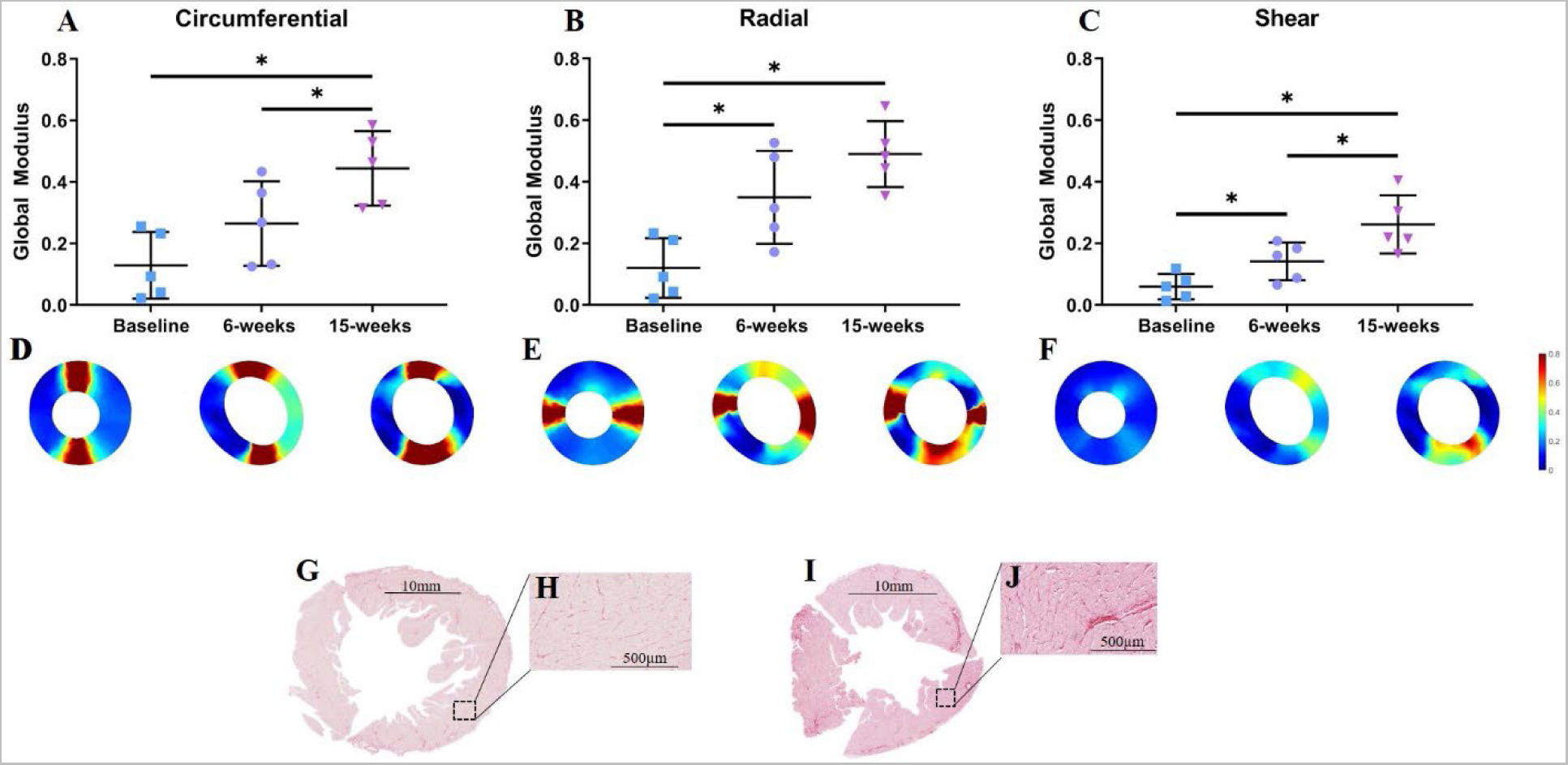
Average global circumferential (A), radial (B), and shear modulus (C) at baseline, 6-weeks, and 15-weeks after doxorubicin administration. Whiskers indicate standard deviation. * indicates *p*<0.05, ** indicates *p* <0.01, and *** indicates *p*<0.001. Representative spatial elasticity maps of circumferential (D), Radial (E), and Shear (F) modulus at each image acquisition. Picrosirius red stained LV of control sample(G,H) and sample at 15-weeks (I,J) indicating increased collagen deposition with corresponding scale bars.

### Segmental Analysis

The LV was partitioned into three segments (septal, posterior, and anterior regions) for segmental analysis of increases in elasticity between image acquisitions. Estimated elastic modulus values were averaged within each segment to determine segmental changes in modulus over time due to doxorubicin administration using two-way ANOVA, shown in Figure 2A-C. The anterior region exhibited a significant increase in all modulus directions between baseline and 15-weeks and was significantly elevated at 15-weeks compared to the septal region in the circumferential and shear directions. Circumferential and shear modulus mimic global analysis and significantly increases from 6-weeks to 15-weeks in the anterior region. Circumferential modulus in the posterior region significantly increased from 6- to 15-weeks. Segmental analysis complements global modulus analysis and shows overall progressive stiffening of the LV due to anthracycline administration.

**Figure 2:**
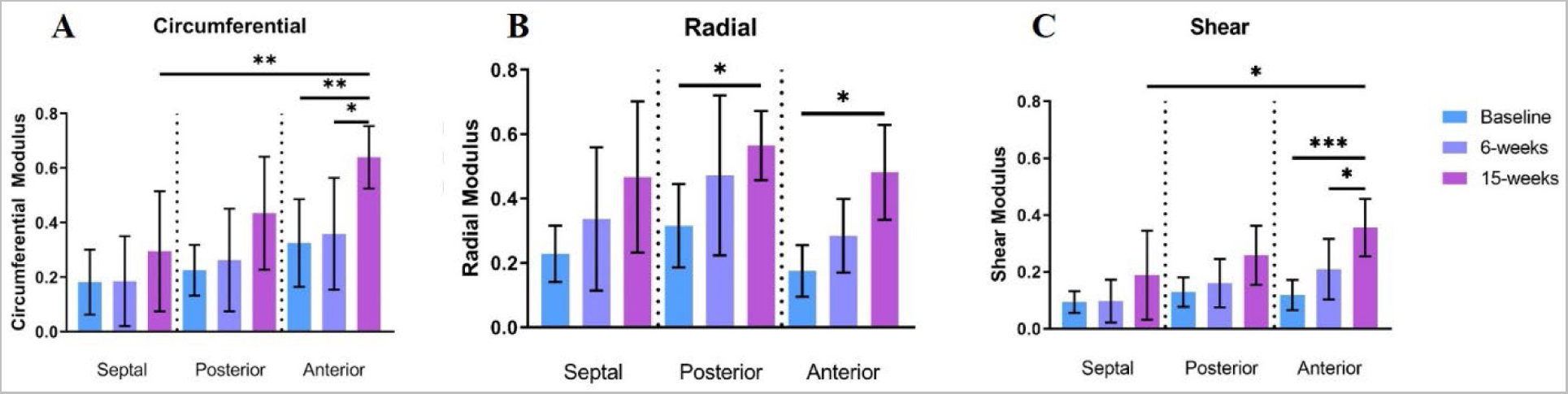
Average regional circumferential (A), radial (B), and shear modulus (C) at baseline, 6-weeks, and 15-weeks after doxorubicin administration. Whiskers indicate standard deviation. * indicates *p*<0.05, ** indicates *p* <0.01, and *** indicates *p*<0.001.

### Correlation between tissue elasticity and pathology

Average global elasticity modulus values were correlated with CVF and cardiomyocyte area to determine elasticity association with histopathology features of cardiotoxicity. Elastic modulus values calculated from baseline images (baseline) were compared to age-matched tissue bank pathological samples (control), and modulus values calculated from images acquired at 15-weeks were directly compared to corresponding pathological data from necropsy at 15-weeks (15-weeks) using a paired analysis pairing age-matched control samples to age-matched experimental samples. Circumferential, radial, and shear modulus increased with increases in CVF and cardiomyocyte area, as shown in Figure 3. Results indicate strong-to-very-strong correlations between global LV circumferential, radial, and shear modulus and both CVF and cardiomyocyte area, with strongest correlation between circumferential and radial modulus with cardiomyocyte area indicated by calculated *p*-values and corresponding Pearson’s correlation coefficients (r = 0.83 - 0.90 for all).

**Figure 3:**
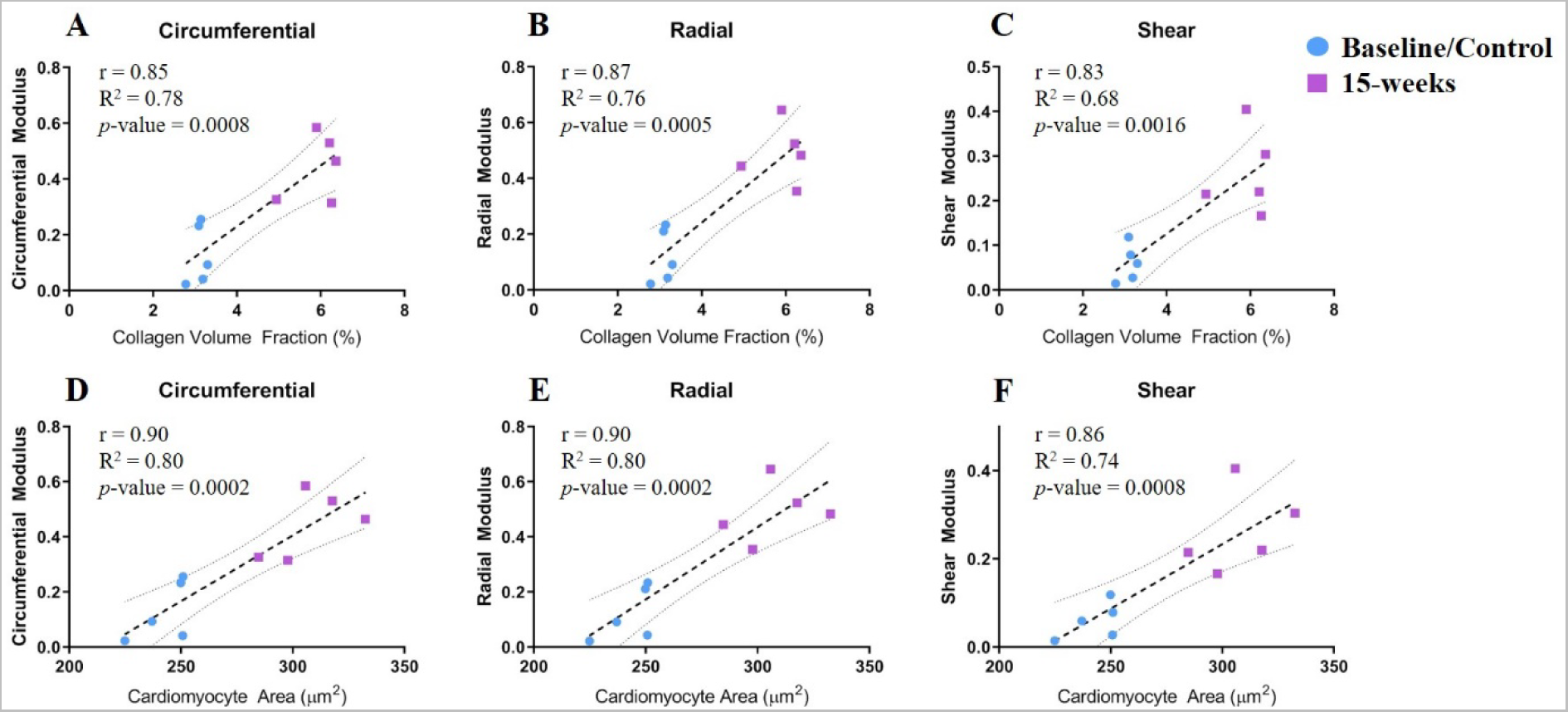
Collagen volume fraction (CVF)(A-C) and cardiomyocyte area (D-F) of age-matched control samples (baseline) and samples acquired at 15-weeks (15-weeks) plotted against circumferential, radial, and shear modulus calculated from baseline and 15-week (15-weeks) imaging data. Dashed line represents linear correlation, with dotted lines representing 95% confidence interval. r, R^2^ and *p*-values for Pearson’s correlation analysis is indicated on each graph.

### Strain Imaging

To assess changes in strain between image acquisition time points, peak circumferential strain at each time point was calculated and is shown in Figure 4A with statistical changes in peak circumferential strain between baseline, 6-weeks, and 15-weeks assessed by t-tests and significance indicated by asterisks. Peak circumferential strain was found to significantly decrease between both baseline and 6-weeks and between baseline and 15-weeks. However, global peak circumferential strain was not significantly different between 6-weeks and 15-weeks, with a trend towards recovery (Figure 4).

**Figure 4:**
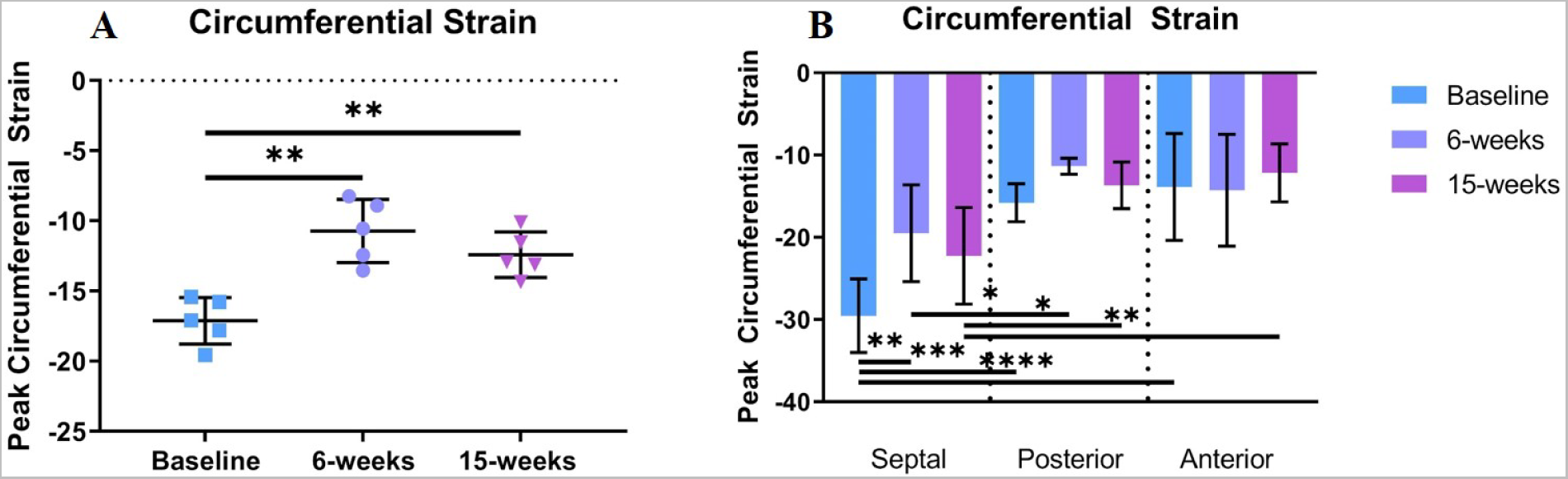
Average global (A) and regional (B) peak circumferential strain, baseline, 6-weeks, and 15-weeks after doxorubicin administration. Whiskers indicate standard deviation. * indicates *p*<0.05, ** indicates *p* <0.01, and *** indicates *p*<0.001.

Peak circumferential strain was also assessed in each segment at each time point, and significant differences between image time point and region was calculated using two-way ANOVA, shown in Figure 4B. Peak circumferential strain did not significantly decrease from baseline to 15-weeks in any LV region. Circumferential strain decreased significantly between baseline and 6-weeks in the septal region. Peak circumferential strain did significantly differ regionally. We also assessed correlation between peak circumferential strain and tissue pathology. Correlations between peak circumferential strain and CVF and cardiomyocyte area are shown in Figure 5. Circumferential strain showed strong correlation with both CVF and cardiomyocyte area.

**Figure 5:**
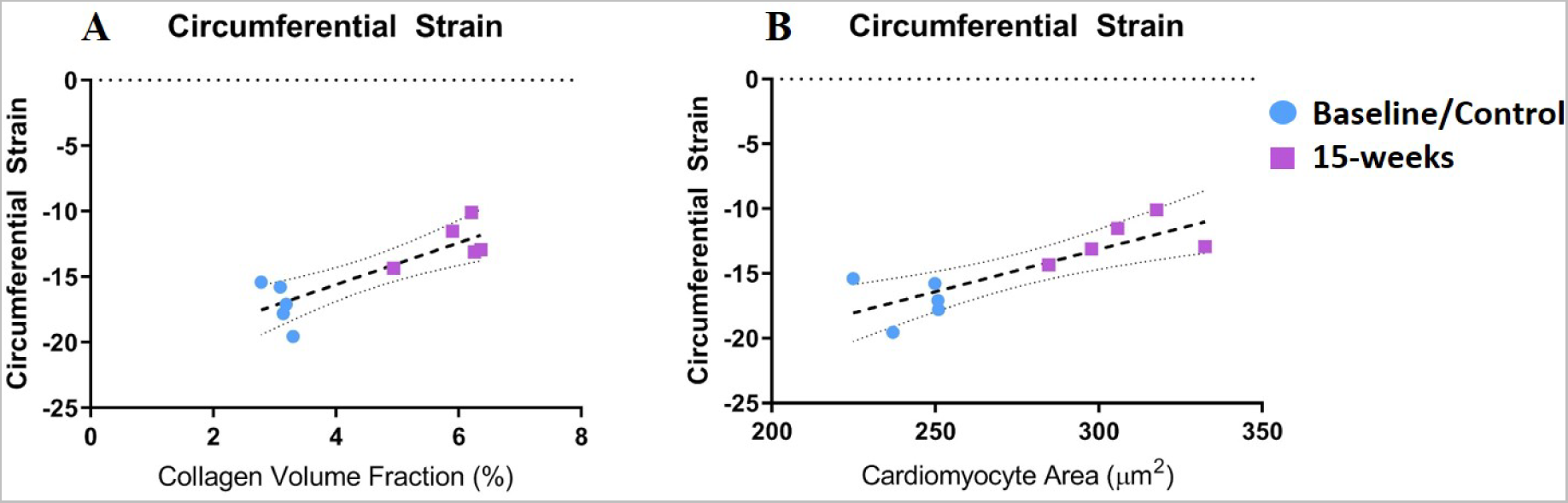
Collagen volume fraction (CVF) (A) and cardiomyocyte area (B) of age-matched control samples (baseline) and samples acquired at 15-weeks (15-weeks) plotted against peak circumferential strain calculated from baseline and 15-week (15-weeks) imaging data.

## Discussion

In this study, we employed BEIM to estimate LV mechanical elasticity on previously-acquired CMR images from African Green monkeys with significant LV dysfunction induced by cardiotoxic anthracycline treatment and assessed their association with histopathological remodeling. We demonstrate that: 1) LV elasticity progressively increases after Dox treatment, and 2) LV elasticity significantly correlates with histopathological markers of cardiotoxicity.

We employed a robust non-human primate model of heart failure induced by doxorubicin treatment. Animals exhibited cardiotoxicity, indicated by a significant decrease in LVEF and CO (> 10% form baseline) and at a histopathological level, Dox induced cardiomyocyte apoptosis accompanied by hypertrophy of the remaining cardiomyocytes and progressive interstitial fibrosis ^16, 32–37^. These cellular and molecular underlying mechanisms of cardiotoxicity potentially compromise cardiac compliance and increase myocardial elasticity, which can be leveraged as a marker of subclinical cardiac injury.

BEIM-assessed LV elasticity metrics of global circumferential, radial, and shear modulus significantly increased from baseline to 15-weeks (Figure 1). We also found a significant change in circumferential and shear modulus between 6-weeks and 15-weeks, and in radial and shear modulus between baseline and 6-weeks. Combined, results indicate progressive increases in myocardial elasticity that occur as early as 6-weeks after the final dose of Dox, with progressive stiffening up to 15-weeks. Progressive stiffening is hypothesized to be a result of increased collagen deposition and cardiomyocyte loss due to anthracycline-induced cardiotoxicity^37, 38^ and is supported by histopathological staining indicating an increase in collagen volume fraction from control samples compared to tissue samples at 15-weeks (Figure 1 G-J). As a result, LV elasticity changes estimated using BEIM may offer a non-invasive strategy for assessing changes in LV collagen deposition indicative of cardiotoxicity.

Spatial analysis is vital to generate hypotheses about regional dependencies of LV remodeling due to cardiotoxic therapies. To assess spatial changes in elasticity, we calculated average modulus within the septal, posterior, and anterior segments of the LV and found significant changes in elasticity in all regions. Modulus in all directions in the anterior region increased between baseline and 15-weeks, and circumferential and shear anterior modulus was significantly increased at 15-weeks from the septal region. Results indicate that anthracycline-induced cardiotoxicity may result in increased ECM remodeling in free wall regions compared with the septal region, though more studies are needed to determine region specific pathological changes due to Dox. Circumferential and shear modulus significantly increased between 6- and 15-weeks, indicating changes in LV myocardial elasticity may continue to occur after therapy has concluded, and that BEIM estimated values of LV elasticity are sensitive to this change. All spatial regions exhibit a trend of increasing LV elasticity between each image acquisition time point, suggesting LV elasticity increases early and continues to change due to Dox.

We also assessed correlations between elasticity and histopathological indicators of cardiotoxicity to validate the ability of BEIM to indicate pathological LV tissue changes. BEIM-estimated changes in LV elasticity correlate strongly with early pathological tissue changes. Circumferential, radial, and shear modulus were found to strongly correlate with both cardiomyocyte area and CVF, validating the utility of BEIM measures of elasticity as an indicator of cardiotoxicity. Circumferential, radial, and shear modulus correlated very strongly with cardiomyocyte area, shown in Figure 3. Overall, BEIM elasticity measures correlated with different mechanisms of tissue stiffening, including both increases in LV stiffness due to increased collagen deposition and increases in passive cardiomyocyte stiffness due to hypertrophic cardiomyocytes, suggesting our methodology is sensitive to different pathological effects of anti-neoplastic therapy on the myocardium.

We also assessed changes in peak LV circumferential strain due to anthracycline therapy, a clinically-accepted CMR metric to assess cardiac function^9^. For short-axis plane images, circumferential strain is considered a better predictor of heart failure than radial strain^7, 39^, therefore only the peak circumferential strain was assessed in this study. We found peak circumferential strain decreased from baseline to 6-weeks and baseline to 15-weeks, but change was insignificant between 6-weeks and 15-weeks. Data is suggestive of strain recovery between 6- and 15-weeks, but strain at 6-weeks may be artificially influenced by hypovolemia, similar to trends seen in LVEF data, as strain has been previously shown to be volume dependent^13, 40^. In contrast, our elasticity estimations show progressive stiffening between these time points, potentially indicating a more sensitive indicator of progressive stiffening than strain imaging. We also assessed regional peak circumferential strain and found significant decline between baseline and 6-weeks only in the septal region. Peak circumferential strain was also found to strongly correlate with cardiomyocyte area and CVF (Figure 4), though correlations were slightly less than elasticity. Note that to facilitate direct comparisons to histopathological tissue that was processed in the short-axis orientation, in this study we only assessed CMR data acquired in the short-axis plane and therefore did not assess elasticity or strain in the longitudinal direction. It is important to note however that CMR circumferential strain has been previously shown to be associated with concurrent CTRCD for identifying cardiotoxicity in breast cancer^6^.

Our method shows the potential to identify subclinical cardiotoxicity by assessing LV elasticity changes, but further studies are needed to address several limitations. In a first-order approximation, the current implementation of our method utilizes an underlying biomechanical model assumption of linear transverse isotropic elasticity. However, nonlinear, anisotropic, viscoelastic behavior due to cardiac geometry and fiber alignment represents a more complex biomechanical model. Intentional simplifications in our model were made to develop a method capable of utilizing routine cardiac imaging for increased clinical translational applicability. To minimize the effects of viscoelastic behavior, we calculate LV elasticity based on deformation during passive diastole in which ventricular pressure changes are minimal. Future work is needed to further account for complex material behavior of the heart. Specialized imaging sequences to indicate cardiac fiber alignment would be needed in addition to routine clinical imaging to define a more rigorous nonlinear anisotropic cardiac behavior. While this would allow for a more accurate representation of cardiac deformation for subsequent elasticity reconstruction, it may potentially limit utility in settings without available advanced imaging data. With respect to study limitations, AGM subjects were used to recapitulate the effect of doxorubicin-induced cardiotoxicity with pathological tissue data available for comparison at study endpoint. However, the use of AGM subjects to obtain LV metrics that were more biosimilar to human subjects, and imaging data obtained through clinically relevant MR sequences resulted in the trade-off of limited sample size. Additionally, age-matched tissue bank samples were used to provide pathological tissue samples that represented baseline measurements to compare to AGM subjects undergoing doxorubicin therapy. We compared baseline LV elasticity results to gender- and age-matched tissue bank samples due to ethical considerations to limit the sacrifice of healthy AGM subjects. While tissue bank samples were acquired from subjects from differing studies, they are representative of healthy AGM subjects at baseline and provide important comparator pathological data.

## Conclusions

In conclusion, this study demonstrates BEIM-estimated elasticity maps identify progressive changes in elasticity and correlates with histopathological tissue changes due to cardiotoxicity in an AGM model of doxorubicin-induced cardiotoxicity. Significant changes in LVEF and CO indicated severe cardiotoxicity due to doxorubicin in this experimental model with histopathological analysis indicating significant changes in cardiomyocyte area, cardiomyocyte number, and CVF. BEIM was used to assess global and regional changes in LV elasticity and results show the potential for BEIM to identify progressive LV stiffening resulting from cardiotoxic effects of anthracycline-based therapy. LV elasticity estimated using BEIM demonstrated strong correlations to cardiomyocyte area and CVF, known to be involved in modulating passive myocardial tissue compliance. Correlations show the potential for LV elasticity measurements to indicate changes in cardiac tissue due to anthracycline-induced cardiotoxicity. As BEIM calculates a different, more functional and direct measurement of elasticity than strain imaging, this preliminary study suggests that BEIM could provide an additional measurement of LV elasticity utilizing clinically relevant imaging and provides significant promise for further interrogation in future studies for identifying subclinical cardiac degradation. Utilizing BEIM to show sensitive spatial LV stiffening based on CMR images acquired in a setting that closely recapitulates clinical workflow highlights the advantages of our methodology for assessing myocardial elasticity changes and potential for this methodology to complement current standard metrics for diagnosing cardiotoxicity. Myocardial elasticity may be an important biomarker for cardiotoxicity as well as other etiologies of cardiac disease. This study provides incentive for future work utilizing BEIM to identify early cardiac disease.

## List of abbreviations

CMR: Cardiac Magnetic Resonance
BEIM: biomechanical model-based elasticity imaging methodology
LV: left ventricle
LVEF: left ventricle ejection fraction
CO: cardiac output
GLS: global longitudinal strain
Dox: doxorubicin
AGM: African Green Monkey
CVF: collagen volume fraction
MR: magnetic resonance

## Declarations

### Ethics approval and consent to participate

This study was approved by the Wake Forest University Institutional Animal Care and Use Committee.

### Consent for publication

Not applicable.

### Availability of data and materials

The datasets used and/or analyzed during the current study are available from the corresponding author on reasonable request.

## Funding

This work is supported by the Williams Family Chair in Breast Oncology and the National Institutes of Health-National Cancer Institute: K25CA204599, P30CA012197.

### Authors’ Contributions

C.M., J.W. and G.M. designed studies. C.M. and S.F. analyzed data. G.M., provided data. C.M., J.W., and G.M. wrote the manuscript text. All authors revised and reviewed the manuscript.

## Acknowledgements

Not applicable.

## References

1. Sturgeon KM, Deng L, Bluethmann SM, Zhou S, Trifiletti DM, Jiang C, Kelly SP, Zaorsky NG. A population-based study of cardiovascular disease mortality risk in US cancer patients. Eur Heart J. 2019;40:3889–3897. doi: 10.1093/eurheartj/ehz766

2. Zhang X, Pawlikowski M, Olivo-Marston S, Williams KP, Bower JK, Felix AS. Ten-year cardiovascular risk among cancer survivors: The National Health and Nutrition Examination Survey. PLoS One. 2021;16:e0247919. doi: 10.1371/journal.pone.0247919

3. Cocco LD, Chiaparini AF, Saffi MAL, Leiria TLL. Global Longitudinal Strain for the Early Detection of Chemotherapy-Induced Cardiotoxicity: A Systematic Review and Meta-analysis. Clin Oncol (R Coll Radiol). 2022;34:514–525. doi: 10.1016/j.clon.2022.05.001

4. Hare JL, Brown JK, Leano R, Jenkins C, Woodward N, Marwick TH. Use of myocardial deformation imaging to detect preclinical myocardial dysfunction before conventional measures in patients undergoing breast cancer treatment with trastuzumab. Am Heart J. 2009;158:294–301. doi: 10.1016/j.ahj.2009.05.031

5. Oikonomou EK, Kokkinidis DG, Kampaktsis PN, Amir EA, Marwick TH, Gupta D, Thavendiranathan P. Assessment of Prognostic Value of Left Ventricular Global Longitudinal Strain for Early Prediction of Chemotherapy-Induced Cardiotoxicity: A Systematic Review and Meta-analysis. JAMA Cardiol. 2019;4:1007–1018. doi: 10.1001/jamacardio.2019.2952

6. Houbois CP, Nolan M, Somerset E, Shalmon T, Esmaeilzadeh M, Lamacie MM, Amir E, Brezden-Masley C, Koch CA, Thevakumaran Y, et al. Serial Cardiovascular Magnetic Resonance Strain Measurements to Identify Cardiotoxicity in Breast Cancer: Comparison With Echocardiography. JACC Cardiovasc Imaging. 2021;14:962–974. doi: 10.1016/j.jcmg.2020.09.039

7. Kim HY, Park SJ, Lee SC, Chang SY, Kim EK, Chang SA, Choi JO, Park SW, Kim SM, Choe YH, et al. Comparison of global and regional myocardial strains in patients with heart failure with a preserved ejection fraction vs hypertension vs age-matched control. Cardiovasc Ultrasound. 2020;18:44. doi: 10.1186/s12947-020-00223-0

8. Pryds K, Larsen AH, Hansen MS, Grondal AYK, Tougaard RS, Hansson NH, Clemmensen TS, Logstrup BB, Wiggers H, Kim WY, et al. Myocardial strain assessed by feature tracking cardiac magnetic resonance in patients with a variety of cardiovascular diseases - A comparison with echocardiography. Sci Rep. 2019;9:11296. doi: 10.1038/s41598-019-47775-4

9. Thavendiranathan P, Poulin F, Lim KD, Plana JC, Woo A, Marwick TH. Use of myocardial strain imaging by echocardiography for the early detection of cardiotoxicity in patients during and after cancer chemotherapy: a systematic review. J Am Coll Cardiol. 2014;63:2751–2768. doi: 10.1016/j.jacc.2014.01.073

10. Guccione JM, McCulloch AD, Waldman LK. Passive material properties of intact ventricular myocardium determined from a cylindrical model. J Biomech Eng. 1991;113:42–55. doi: 10.1115/1.2894084

11. Jordan JH, Castellino SM, Melendez GC, Klepin HD, Ellis LR, Lamar Z, Vasu S, Kitzman DW, Ntim WO, Brubaker PH, et al. Left Ventricular Mass Change After Anthracycline Chemotherapy. Circ Heart Fail. 2018;11:e004560. doi: 10.1161/CIRCHEARTFAILURE.117.004560

12. Jefferies JL, Mazur WM, Howell CR, Plana JC, Ness KK, Li Z, Joshi VM, Green DM, Mulrooney DA, Towbin JA, et al. Cardiac remodeling after anthracycline and radiotherapy exposure in adult survivors of childhood cancer: A report from the St Jude Lifetime Cohort Study. Cancer. 2021;127:4646–4655. doi: 10.1002/cncr.33860

13. Jordan JH, Sukpraphrute B, Melendez GC, Jolly MP, D’Agostino RB, Jr., Hundley WG. Early Myocardial Strain Changes During Potentially Cardiotoxic Chemotherapy May Occur as a Result of Reductions in Left Ventricular End-Diastolic Volume: The Need to Interpret Left Ventricular Strain With Volumes. Circulation. 2017;135:2575–2577. doi: 10.1161/CIRCULATIONAHA.117.027930

14. Miller CE, Jordan JH, Thomas A, Weis JA. Developing a biomechanical model-based elasticity imaging method for assessing hormone receptor positive breast cancer treatment-related myocardial stiffness changes. J Med Imaging (Bellingham). 2021;8:056002. doi: 10.1117/1.JMI.8.5.056002

15. Miller CE, Jordan JH, Douglas E, Ansley K, Thomas A, Weis JA. Reproducibility assessment of a biomechanical model-based elasticity imaging method for identifying changes in left ventricular mechanical stiffness. J Med Imaging (Bellingham). 2022;9:056001. doi: 10.1117/1.JMI.9.5.056001

16. Melendez GC, Vasu S, Lesnefsky EJ, Kaplan JR, Appt S, D’Agostino RB, Jr., Hundley WG, Jordan JH. Myocardial Extracellular and Cardiomyocyte Volume Expand After Doxorubicin Treatment Similar to Adjuvant Breast Cancer Therapy. JACC Cardiovasc Imaging. 2020;13:1084–1085. doi: 10.1016/j.jcmg.2019.10.020

17. Levick SP, Soto-Pantoja DR, Bi J, Hundley WG, Widiapradja A, Manteufel EJ, Bradshaw TW, Melendez GC. Doxorubicin-Induced Myocardial Fibrosis Involves the Neurokinin-1 Receptor and Direct Effects on Cardiac Fibroblasts. Heart Lung Circ. 2019;28:1598–1605. doi: 10.1016/j.hlc.2018.08.003

18. Melendez GC, Li J, Law BA, Janicki JS, Supowit SC, Levick SP. Substance P induces adverse myocardial remodelling via a mechanism involving cardiac mast cells. Cardiovasc Res. 2011;92:420–429. doi: 10.1093/cvr/cvr244

19. Melendez GC, McLarty JL, Levick SP, Du Y, Janicki JS, Brower GL. Interleukin 6 mediates myocardial fibrosis, concentric hypertrophy, and diastolic dysfunction in rats. Hypertension. 2010;56:225–231. doi: 10.1161/HYPERTENSIONAHA.109.148635

20. Heiberg E, Sjogren J, Ugander M, Carlsson M, Engblom H, Arheden H. Design and validation of Segment--freely available software for cardiovascular image analysis. BMC Med Imaging. 2010;10:1. doi: 10.1186/1471-2342-10-1

21. Sechtem U, Pflugfelder PW, Gould RG, Cassidy MM, Higgins CB. Measurement of right and left ventricular volumes in healthy individuals with cine MR imaging. Radiology. 1987;163:697–702. doi: 10.1148/radiology.163.3.3575717

22. Drafts BC, Twomley KM, D’Agostino R, Jr., Lawrence J, Avis N, Ellis LR, Thohan V, Jordan J, Melin SA, Torti FM, et al. Low to moderate dose anthracycline-based chemotherapy is associated with early noninvasive imaging evidence of subclinical cardiovascular disease. JACC Cardiovasc Imaging. 2013;6:877–885. doi: 10.1016/j.jcmg.2012.11.017

23. Thirion JP. Image matching as a diffusion process: an analogy with Maxwell’s demons. Med Image Anal. 1998;2:243–260. doi: 10.1016/s1361-8415(98)80022-4

24. Yoo TS, Ackerman MJ, Lorensen WE, Schroeder W, Chalana V, Aylward S, Metaxas D, Whitaker R. Engineering and algorithm design for an image processing Api: a technical report on ITK--the Insight Toolkit. Stud Health Technol Inform. 2002;85:586–592.

25. McCormick M, Liu X, Jomier J, Marion C, Ibanez L. ITK: enabling reproducible research and open science. Front Neuroinform. 2014;8:13. doi: 10.3389/fninf.2014.00013

26. Liu DC, Nocedal J. On the Limited Memory Bfgs Method for Large-Scale Optimization. Math Program. 1989;45:503–528. doi: Doi 10.1007/Bf01589116

27. Nocedal J. Updating Quasi-Newton Matrices with Limited Storage. Mathematics of Computation. 1980;35:773–782. doi: Doi 10.2307/2006193

28. Sohn DW, Lee SP, Kim HK, Kim YJ, Yoo BW, Kim HC. LV peak instantaneous wall stress versus time-stress-integral as measures of afterload in aortic stenosis. Heart. 2015;101:478–483. doi: 10.1136/heartjnl-2014-307151

29. Cerqueira MD, Weissman NJ, Dilsizian V, Jacobs AK, Kaul S, Laskey WK, Pennell DJ, Rumberger JA, Ryan T, Verani MS, et al. Standardized myocardial segmentation and nomenclature for tomographic imaging of the heart. A statement for healthcare professionals from the Cardiac Imaging Committee of the Council on Clinical Cardiology of the American Heart Association. Circulation. 2002;105:539–542. doi: 10.1161/hc0402.102975

30. Heyde B, Jasaityte R, Barbosa D, Robesyn V, Bouchez S, Wouters P, Maes F, Claus P, D’Hooge J. Elastic image registration versus speckle tracking for 2-D myocardial motion estimation: a direct comparison in vivo. IEEE Trans Med Imaging. 2013;32:449–459. doi: 10.1109/TMI.2012.2230114

31. Morais P, Heyde B, Barbosa D, Queirós S, Claus P, D’hooge J. Cardiac motion and deformation estimation from tagged MRI sequences using a temporal coherent image registration framework. Paper/Poster presented at: Functional Imaging and Modeling of the Heart: 7th International Conference, FIMH 2013, London, UK, June 20-22, 2013. Proceedings 7; 2013;

32. Vejpongsa P, Yeh ET. Prevention of anthracycline-induced cardiotoxicity: challenges and opportunities. J Am Coll Cardiol. 2014;64:938–945. doi: 10.1016/j.jacc.2014.06.1167

33. Cai F, Luis MAF, Lin X, Wang M, Cai L, Cen C, Biskup E. Anthracycline-induced cardiotoxicity in the chemotherapy treatment of breast cancer: Preventive strategies and treatment. Mol Clin Oncol. 2019;11:15–23. doi: 10.3892/mco.2019.1854

34. Zeiss CJ, Gatti DM, Toro-Salazar O, Davis C, Lutz CM, Spinale F, Stearns T, Furtado MB, Churchill GA. Doxorubicin-Induced Cardiotoxicity in Collaborative Cross (CC) Mice Recapitulates Individual Cardiotoxicity in Humans. *G3* (*Bethesda*). 2019;9:2637-2646. doi: 10.1534/g3.119.400232

35. Ghigo A, Li M, Hirsch E. New signal transduction paradigms in anthracycline-induced cardiotoxicity. Biochim Biophys Acta. 2016;1863:1916–1925. doi: 10.1016/j.bbamcr.2016.01.021

36. Karagiannis TC, Lin AJ, Ververis K, Chang L, Tang MM, Okabe J, El-Osta A. Trichostatin A accentuates doxorubicin-induced hypertrophy in cardiac myocytes. Aging (Albany NY). 2010;2:659–668. doi: 10.18632/aging.100203

37. Lushnikova EL, Klinnikova MG, Molodykh OP, Nepomnyashchikh LM. Morphological manifestations of heart remodeling in anthracycline-induced dilated cardiomyopathy. Bull Exp Biol Med. 2004;138:607–612. doi: 10.1007/s10517-005-0138-0

38. Unverferth DV, Baker PB, Pearce LI, Lautman J, Roberts WC. Regional myocyte hypertrophy and increased interstitial myocardial fibrosis in hypertrophic cardiomyopathy. Am J Cardiol. 1987;59:932–936. doi: 10.1016/0002-9149(87)91128-3

39. Buss SJ, Krautz B, Hofmann N, Sander Y, Rust L, Giusca S, Galuschky C, Seitz S, Giannitsis E, Pleger S, et al. Prediction of functional recovery by cardiac magnetic resonance feature tracking imaging in first time ST-elevation myocardial infarction. Comparison to infarct size and transmurality by late gadolinium enhancement. Int J Cardiol. 2015;183:162–170. doi: 10.1016/j.ijcard.2015.01.022

40. Melendez GC, Sukpraphrute B, D’Agostino RB, Jr., Jordan JH, Klepin HD, Ellis L, Lamar Z, Vasu S, Lesser G, Burke GL, et al. Frequency of Left Ventricular End-Diastolic Volume-Mediated Declines in Ejection Fraction in Patients Receiving Potentially Cardiotoxic Cancer Treatment. Am J Cardiol. 2017;119:1637–1642. doi: 10.1016/j.amjcard.2017.02.008

